# Semi-automated analysis of an optical ATP indicator in neurons

**DOI:** 10.1101/2021.09.20.461141

**Authors:** Taher Dehkharghanian, Arsalan Hashemiaghdam, Ghazaleh Ashrafi

**Author notes:** These authors contributed equally to this manuscript (Taher Dehkharghanian, Arsalan Hashemiaghdam).

## Abstract

**Significance:** The firefly enzyme luciferase has been used in a wide range of biological assays, including bioluminescence imaging of ATP. The biosensor Syn-ATP utilizes subcellular targeting of luciferase to nerve terminals for optical measurement of ATP in this compartment. Manual analysis of Syn-ATP signals is challenging due to signal heterogeneity and cellular motion in long imaging sessions. Here, we have leveraged machine learning tools to develop a method for analysis of bioluminescence images.

**Aim:** Our goal was to create a semi-automated pipeline for analysis of bioluminescence imaging to improve measurements of ATP content in nerve terminals.

**Approach:** We developed an image analysis pipeline that applies machine learning toolkits to distinguish neurons from background signals, and excludes neural cell bodies, while also incorporating user input.

**Results:** Side-by-side comparison of manual and semi-automated image analysis demonstrated that the latter improves precision and accuracy of ATP measurements.

**Conclusions:** Our method streamlines data analysis and reduces user-introduced bias, thus enhancing the reproducibility and reliability of quantitative ATP imaging in nerve terminals.

## Introduction

In nature, living organisms as diverse as bacteria, fireflies, copepods, and sea pansies produce natural light through bioluminescence ^1^. These organisms express the enzyme luciferase that emits visible light when it catalyzes the oxidation of its substrate luciferin powered by adenosine triphosphate (ATP). Bioluminescence imaging is a sensitive technique that relies on the detection of light emitted from the luciferase reaction ^2^. At saturating luciferin concentrations, luciferase light emission is proportional to ATP level, thus luciferase can be used as a sensitive cellular ATP sensor. Similar to fluorescence, electrons are excited to a higher energy level and emit photons as they return to their resting level ^3^. However, the excitation energy in bioluminesce is provided by the chemical reaction rather than exogenous illumination as in fluorescence (**Fig. 1A**). As a result, bioluminesce does not suffer from photobleaching of excited molecules or phototoxicity. Another advantage of bioluminescence as compared to fluorescence is lower background which allows for higher sensitivity and improved signal to noise ratio (SNR). In fluorescence imaging, biological materials emit significant endogenous fluorescence signals (autofluorescence), particularly in the green emission range, while autoluminescence of most cells is very low ^4^. As such, bioluminescence imaging, despite dimmer signals, is ideally suited for sensitive assays of biological activity.

**Figure 1.**
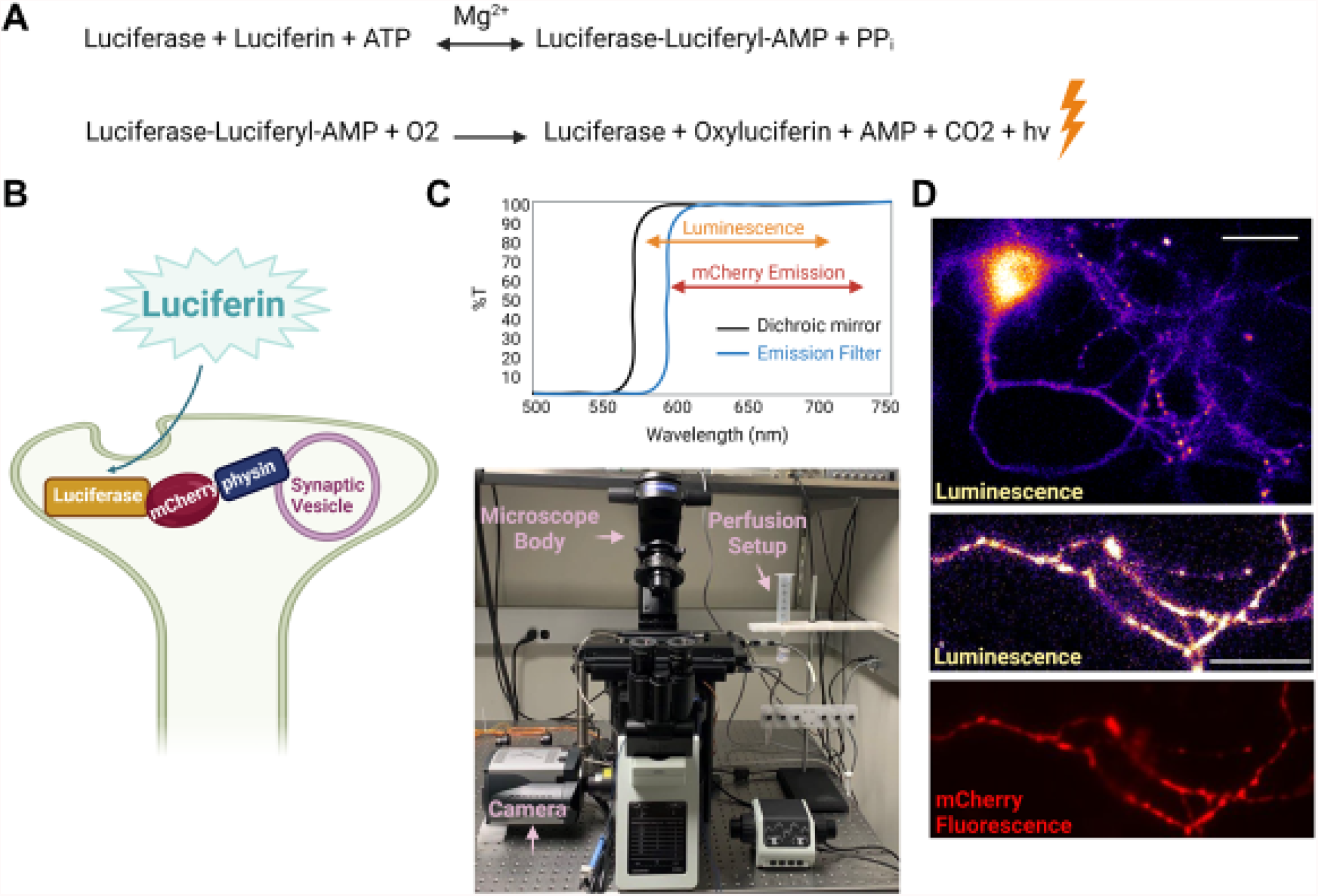
Bioluminescence imaging of cytosolic ATP in nerve terminals. **A)** The bioluminescence chemical reaction in which the enzyme luciferase uses luciferin and ATP to produce light denoted as hν. **B)** Schematic of a hippocampal nerve terminal expressing Syn-ATP in which luciferase is anchored to synaptic vesicles and mCherry is used as an inert fluorophore.**C)** An optimized dual fluorescence and luminescence microscopy setup (bottom) where a long-pass 590 nm filter replaces an emission filter to maximize luminescence photon collection (top). **D)** Representative luminescence and mCherry fluorescence images of a hippocampal neuron (top) and an axon bearing several nerve terminals (bottom). scale bar, 30 µm.

The first practical application of bioluminescence was the development of a reporter for gene expression using the North American firefly (*Photinus pyralis*) luciferase, which emits yellow-green light (emission peak at 557 nm) ^5,6^. Since then, novel mutant bioluminescent reporters emitting red light (> 600 nm) have been engineered to improve tissue penetration for *in vivo* bioluminescence imaging ^7^. Further modifications to luciferase thermostability and catalytic activity have led to the development of ATP sensors for monitoring subcellular ATP levels ^8^. Capitalizing on these improvements, a presynaptic ATP sensor, “Syn-ATP,” was developed to monitor ATP levels in the nerve terminals of cultured hippocampal neurons, particularly to investigate the energetic demands of electrical activity ^9^. Syn-ATP is a genetically encoded optical reporter of ATP, available on Addgene (plasmid # 51819; RRID: Addgene_51819) in which luciferase is targeted to synaptic vesicles through fusion with synaptophysin and additionally tagged with the fluorophore mCherry to normalize for reporter expression level (**Fig. 1B**).

Since its development, imaging data from Syn-ATP assays have been analyzed manually with the software Image J and Microsoft Excel. In this method, the ImageJ Plugin Time Series Analyzer allows for manual selection of regions of interest (ROIs) corresponding to individual nerve terminals, followed by calculation of signal intensity of fluorescence and luminescence over multiple time frames. Raw signal intensities are then subjected to background subtraction. Background-corrected luminescence intensities of individual terminals are then normalized to mCherry fluorescence to correct for variability in Syn-ATP expression and/or changes in the focal plane during imaging. To optimize performance, users need to select the maximal number of mCherry-positive terminals in a recorded field while excluding cell bodies or large cellular clumps. As described above, manual analysis is undesirable because it is time-consuming and subject to user bias. In addition, cellular motion and frame-to-frame movement of the selected ROIs in lengthy (several minutes) experiments complicates data analysis. To address the limitations of manual analysis, we have developed a semi-automated analysis pipeline based on machine-learning algorithms that measures Syn-ATP luminescence signals in individual neurons with appropriate background correction and normalization to fluorescence signals.

## Materials and Methods

All animal experiments were performed with wild-type rats of the Sprague-Dawley strain in accordance with protocols approved by the IACUC at Washington University School of Medicine in St. Louis. Hippocampi were dissected from 0-2 days-old neonatal rats of a mixed (male and female) litter, dissociated, and plated on poly-ornithine coated coverslips as previously described ^10^. Hippocampal neurons at DIV 14-20 were mounted in a laminar flow perfusion chamber, maintained at 37°C with an OkoLab stage top incubator in Tyrode’s buffer containing (in mM) 119 NaCl, 2.5 KCl, 2 CaCl_2_, 2 MgCl_2_, 50 HEPES (pH 7.4), 2 D-Luciferin potassium salt (Gold Biotechnology), 1.25 lactate and 1.25 pyruvate, supplemented with 10 µM 6-cyano-7nitroquinoxalibe-2, 3-dione (CNQX), and 50 µM DL-2-amino-5phosphonovaleric acid (APV) to inhibit postsynaptic responses. Live imaging of the hippocampal neurons was performed on a custom-built, inverted Olympus IX83 epifluorescence microscope equipped for luminescence and fluorescence imaging. Fluorescence excitation of mCherry was achieved with the TTL-controlled Cy3 channel of a Lumencor Aura III light engine. Both mCherry emission and Syn-ATP luminescence were directed through a Chroma 590nm long-pass filter and an Olympus UPlan Fluorite 40X 1.3 NA objective. Image acquisition was performed with an Andor iXon Ultra 897 camera, cooled to -95°C to minimize the camera detection noise. Platinum-Iridium electrodes were used to evoke action potentials with 1ms electrical pulses creating field potentials of ∼10 V/cm. In each experiment, data was collected from at least 10 coverslips from 3 independent cultures prepared from separate litters. Unless otherwise indicated, all chemicals were obtained from Sigma-Aldrich. Image analysis was performed with Image J Time Series Analyzer (manual analysis) or the proposed semi-automated algorithm coded in python programming language, using Jupyter notebook, and scikit-learn machine learning library ^11,12^. Data visualization and statistical analysis were performed in GraphPad Prism v9.0.

## Results

Hippocampal neurons expressing the Syn-ATP sensor were imaged (10 seconds at 2 Hz) for mCherry fluorescence using Cy3 excitation light, followed by a single luminescence frame collected with exposure time of 60 seconds. Emission light from luminescence and fluorescence were directed through a 590nm long-pass filter, instead of a conventional emission filter, to maximize luminescence photon collection (**Fig. 1C**). The alternating acquisition of fluorescence and luminescence images was repeated throughout the experiment to constitute multiple timepoints. The 20-frame fluorescence movie was averaged into a single image, hereafter referred to as the fluorescent image (**Fig. 1D**).

Given that a large and labeled dataset of hippocampal nerve terminals was not publicly available, we decided to implement unsupervised machine learning algorithms for analysis of Syn-ATP images rather than supervised algorithms for detection of individual nerve terminals. Our analysis pipeline is composed of three main steps: (1) background detection, (2) cell body detection, and (3) signal estimation. In order to improve consistency, user input is requested to modify the model’s output for several steps of the process. Our images have two channels (mCherry fluorescence and luminescence) and we used the luminescence channel for background and cell body detection due to its low background signal. We then applied onto fluorescence images the background and cell body masks obtained from the luminescence channel. Details of this semi-automated image analysis pipeline and evaluation of its performance are outlined below.

### Development of a semi-automated analysis pipeline

The first step of the proposed pipeline was to distinguish neurons from the background. The luminescence frame was used as the input image and is downsampled from its original size of 512×512 pixels to 128×128 pixels. Since nerve terminals are about 1.5 µm (∼4 pixels) in diameter, downsampling of the images helps to reduce the likelihood of misidentification of noise as terminals. The resultant image was then divided into regions of 4×4 pixels, which in our microscope setup with 40x objective magnification, corresponds to ∼ 6.4 × 6.4 µm squares. Each region is then represented by its median pixel value.

In the next step, pixel intensities were transformed into a one-dimensional vector of size 16,384 (128×128). K-means clustering from the scikit-learn python library was applied to this vector to produce two clusters, one corresponding to the image background, and the other to the regions of interest ^12,13^. Since K-means is an unsupervised machine learning algorithm, it does not require any training prior to application. We observed that this clustering scheme accurately distinguished background from regions of interest (**Fig. 2A**). We used luminescence images to create a background mask and applied the same mask on fluorescence images (**Fig. 2B**). Background signal in each channel was then calculated as the average intensity of the background pixels.

**Figure 2.**
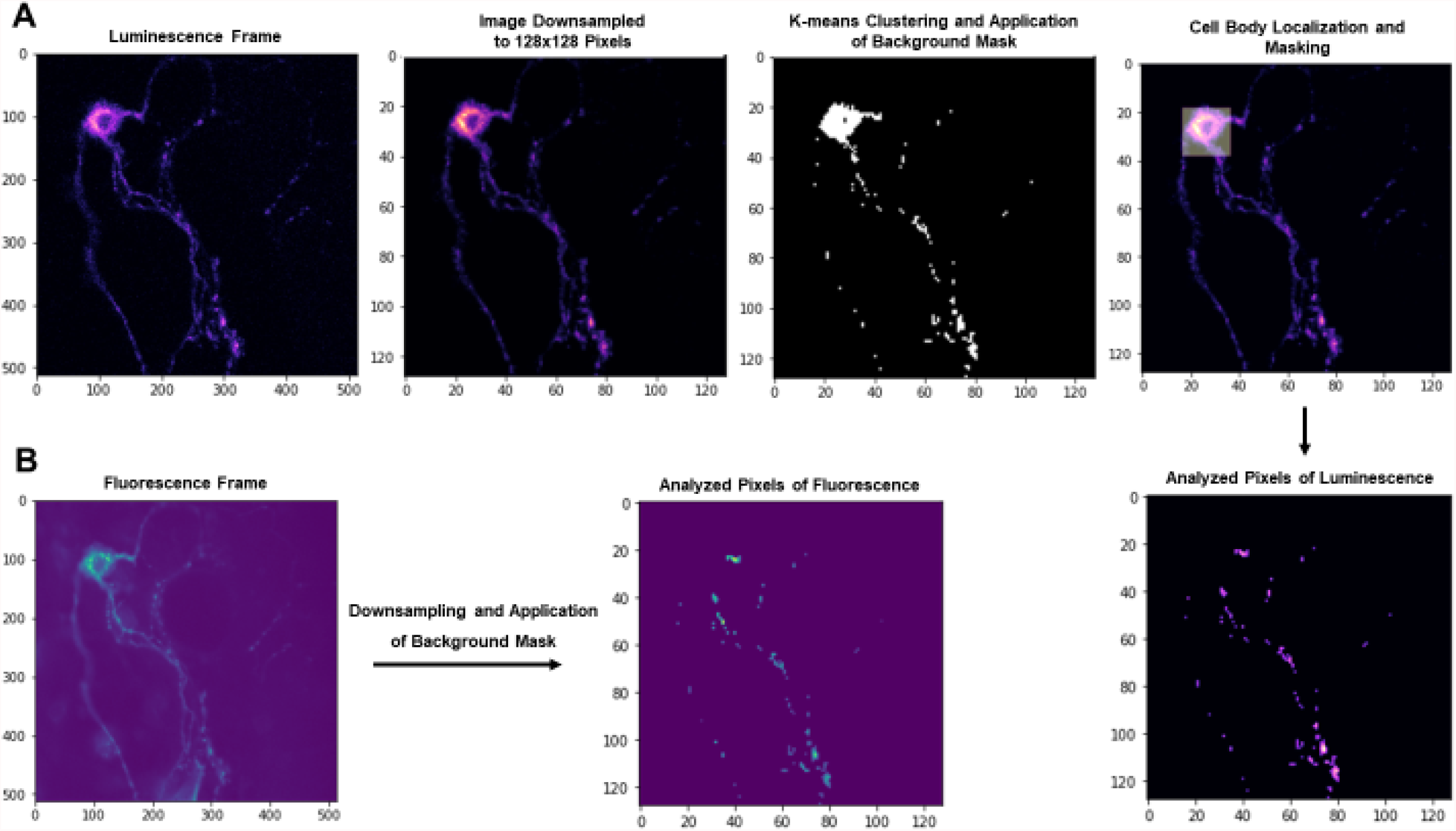
An Image analysis pipeline for background signal determination and cell body removal. **A)** The luminescence image of a neuron was downsampled from 512×512 pixels to 128×128 pixels. K-means clustering algorithm was implemented on pixel values to produce two complementary clusters of background and desired signals. A background mask was applied to remove background signals from the image (black and white panel). Next, the region with the highest total signal intensity was detected and deemed as the cell body. Both background and cell body were removed from further analysis. **B)** Background and cell body masks generated from the luminescence image were applied to the fluorescence image.

Syn-ATP is first synthesized in neural cell bodies where it enters the secretory pathway for trafficking to terminals. However, inclusion of Syn-ATP data from the cell body may be confounding because ATP metabolism in this compartment may differ from nerve terminals. Therefore, the second step in our algorithm was to detect the neural cell body and exclude it from further analysis. In this step, the user provides input to set the custom width (d) of the cell body. Alternatively, the default preset value is set at 32 pixels which corresponds to 50 microns in our setup. A square of size d × d pixels was then moved one pixel at a time, and the mean pixel intensity was calculated for each region. The region with the highest mean intensity was defined as the cell body (**Fig. 2A**). Since neural cell bodies have variable shapes and sizes, and a cell body is not captured in some images, the user is asked to provide their input determining whether the cell body is accurately detected. Otherwise, the user has the option to manually set a custom-sized bounding box to cover the cell body. Once detected, the cell body was masked, as was done for the background, and omitted from further analysis in both luminescence and fluorescence images.

The final step in our pipeline was the determination of signal intensity from masked images. The total intensity of pixels in each of the luminescence and fluorescence images was calculated, followed by subtraction of background intensity to determine net luminescence and fluorescence intensities. This process was iterated for each imaging time point. The L/F value (see below) was plotted against time to represent the ATP content of nerve terminals in a single neuron over time.

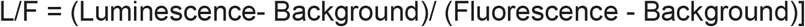

It is important to note that while L/F is directly proportional to ATP concentration in nerve terminals, the absolute L/F value is dependent on image acquisition parameters that may need to be modified during a project. For modifications that produce a linear change in signal intensity, such as adjustments to camera electron-multiplying (EM) gain settings, we introduced a simple true/false argument to adjust calculations of ATP levels by a coefficient factor.

### Performance Evaluation

To evaluate the performance of our analysis pipeline, raw fluorescence and luminescence images from neural samples (n=26 neurons) were analyzed both manually and with our semi-automated tool. The experiment consisted of four baseline 1-minute time points, followed by 1 minute of electrical stimulation at 10 Hz applied between time points 4 and 5. Neurons were imaged for 3 additional time points after stimulation. The reduction in L/F values (**Fig. 3A**) during electrical stimulation was previously attributed to acidification of the cytosol which reduces the enzymatic activity of luciferase given that its pKa (7.03) is close to the cytosolic pKa (6.8). Indeed, both manual and semi-automated analysis methods detect a decline in L/F values during activity which can be fully corrected by taking into account the pH effects using previously described correction factors (**Fig. 3B**) ^9^.

**Figure 3.**
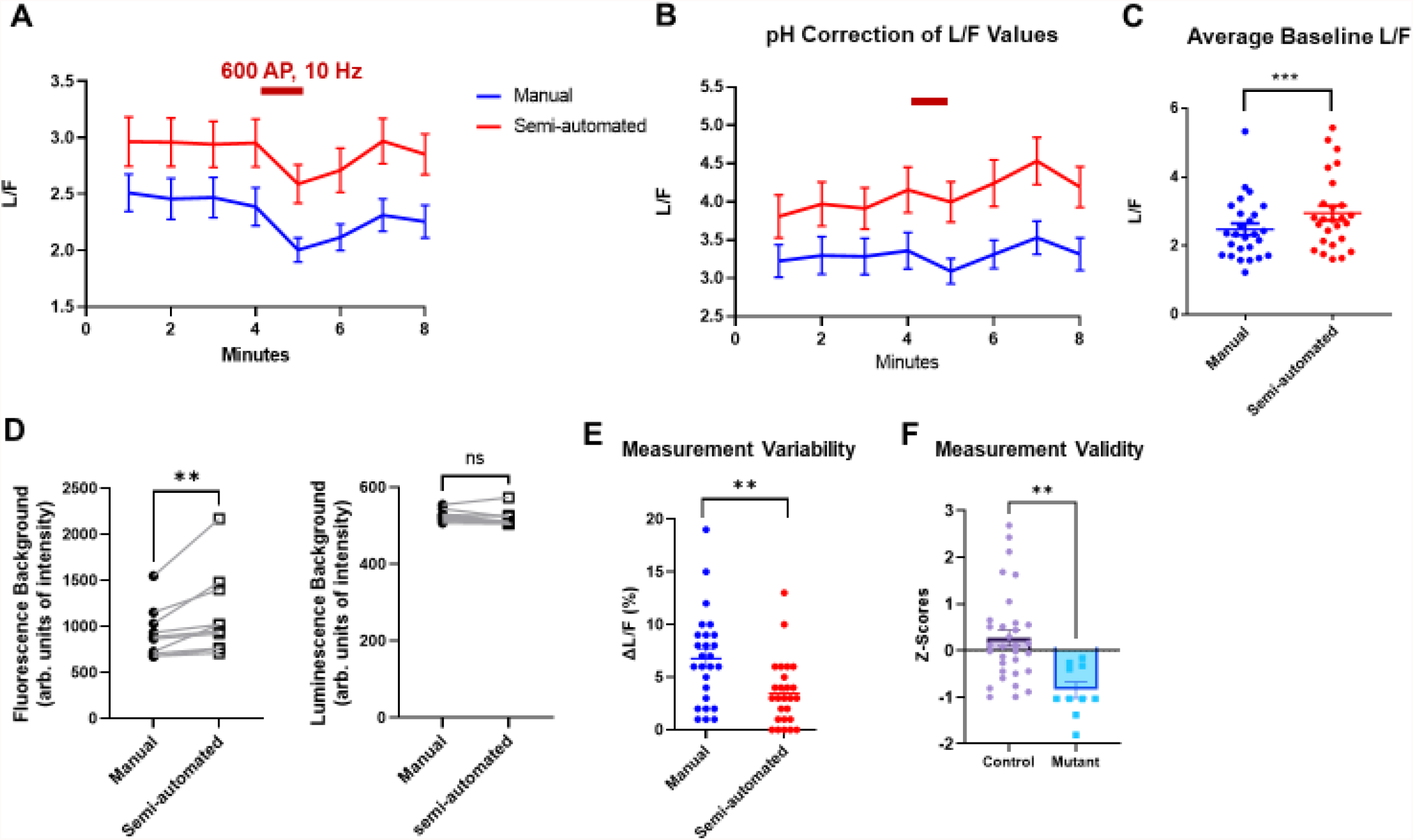
Quantitative comparison of Syn-ATP image analysis by the manual and semi-automated methods. Hippocampal neurons expressing Syn-ATP (n=26) were imaged for 8 minutes, and were electrically stimulated for 1 min at 10Hz frequency (crimson bar). **A)** Average traces of Syn-ATP luminescence normalized by fluorescence intensity (L/F) analyzed by manual and semi-automated methods (n=52 neurons). **B)** L/F traces were corrected for cytosolic pH changes that occur during electrical stimulation. **C)** Baseline pre-stimulation L/F values obtained from semi-automated analysis were significantly higher than manual analysis (paired t-test: p=0.0006). **D)** Semi-automated analysis yielded higher background fluorescence values than manual analysis (paired Wilcoxon test: p=0.002, n=10 neurons) while not affecting luminescence background determination (paired Wilcoxon test: p=0.084, n=10 neurons). **E)** Measurement variability of pre-stimulus L/F values was determined as % deviation from the mean of each neuron (ΔL/F), indicating lower variability with semi-automated analysis (paired t-test: p=0.001, n=52 neurons). **F)** Measurement validity of semi-automated method was assessed by comparing z-scores of two population of control and mitochondrial mutant neurons with different baseline L/F values, indicating significantly lower z-scores for the mutant (Unpaired t-test, p=0.001, ctrl=32 neurons, mutant=10 neurons).

Compared to manual analysis, our semi-automatic method generated higher L/F values (**Fig. 3C**). We speculated this was at least partly due to the more precise determination of background signals through our clustering algorithm. In fact, we examined this hypothesis with a subset of randomly selected neurons from our dataset and demonstrated that background fluorescence values calculated with our semi-automated pipeline were ∼20% higher than manual analysis. The mean intensity for manual and semi-automated were 1120 +/-142 and 921+/- 87 units, respectively. (p-value =0.002) (**Fig. 3D**). In contrast, background luminescence values were not significantly different in the two methods (p-value=0.08) (**Fig. 3D**). The higher fluorescence background values would result in lower background-subtracted F values when using the semi-automated pipeline thus raising the L/F compared to manual measurements as we observed (**Fig. 3C**).

A major challenge in manual analysis of Syn-ATP is the high variability in L/F measurements of the same neuron over time. To determine whether semi-automated yields more consistent values, we compared the variability of L/F measurements obtained from our semi-automated pipeline to manual analysis. In our experiments, the time points prior to stimulation represent baseline ATP levels with minimal biological variation. We calculated the L/F values of the initial three timepoints using both methods and determined the measurement variability (ΔL/F) as % deviation from the mean baseline L/F value for each neuron. The semi-automated approach significantly decreased measurement variabilities (p-value = 0.001) thus increasing the reliability of Syn-ATP analysis. The variation among the three baseline data points across cells (n=26) was 3.4+/-0.9% and 6.8 +/-0.6% in semi-automated and manual analysis, respectively (**Fig. 3E**). These findings demonstrate that our pipeline improves measurement consistency, by minimizing user sampling bias.

We then sought to assess the validity of the semi-automated pipeline in correctly identifying differences in L/F levels in distinct neural populations. Syn-ATP experiments were performed in a population of control neurons and neurons carrying a mitochondrial mutation that impairs mitochondrial ATP production. Following determination of baseline L/F values as in **Fig. 3C**, the two populations were combined, mean and standard deviation of baseline L/F values were calculated, and the z-score of each neuron was determined. Comparison of z-scores for control and mutant neurons revealed significantly lower z-score in the mutant (control: 0.027+/-0.17, mutant: -1.81+/-0.17, p-value=0.001) Therefore, we conclude that the semi-automated method successfully distinguishes differences in Syn-ATP L/F values of distinct genotypes (**Fig. 3F**).

## Discussions and Conclusions

Syn-ATP represents a powerful and robust application of bioluminescence imaging for measurement of ATP levels in nerve terminals. However, analysis of Syn-ATP data has been challenging due to the potential for user selection bias. Standardizing image analysis through advanced computational methods, particularly machine learning, would improve data accuracy and reproducibility. Here, we developed a semi-automated pipeline that facilitates the analysis of dual fluorescence and luminescence images. First, our pipeline enables the user to determine background signals in both channels in an unbiased manner. Second, it enables the user to mask specified regions such as the cell body of a neuron. In imaging sessions that last for several minutes, axonal movement shifts the position of nerve terminals in the field of view and poses challenges for manual tracking of several data points in an image stack. Our approach circumvents this problem by analyzing the entire image rather than individual user-selected ROIs representing nerve terminals.

Despite its advantages, we acknowledge that our semi-automated approach has its own limitations. For example, sudden changes in the background signal interfere with the code and reduce the reliability of the results. Furthermore, user input is required to validate selection of the cell body. Our future endeavors with expanded datasets would be directed toward full automation of the code as well as resolving technical issues that arise from faulty image acquisition.

In summary, we have developed a semi-automated pipeline for analysis of dual fluorescence/luminescence imaging of ATP in nerve terminals. Our pipeline analyzes signals in the entire field of view thereby reducing user sampling error. It also standardizes background signal measurement and reduces variability in measurement of ATP level in nerve terminals.

The code is publicly available on the GitHub library (https://github.com/ashrafilab/SynATP-Analysis) and can be modified for customized bioluminescence image analysis.

## Contributions

T.D. and A.H. share co-first authorship. G.A., A.H., and T.D. conceptualized, designed the method, and wrote the manuscript. T.D. wrote the code in Python. G.A. and A.H. performed experiments and collected the data. G.A. is the corresponding author and supervised the correspondence.

## Disclosure

The authors declare no competing interests.

## Acknowledgments

This study was supported by the McDonnell Center for Cellular and Molecular Neuroscience Small Grants Program and the Esther A. & Joseph Klingenstein Fund. The authors would like to thank Javid Dadashkarimi for proofreading the manuscript, and Marissa Laramie for preparation of neural cultures. Figure 1 was prepared using the BioRender software (Biorender.com).

## Code, Data, and Materials Availability

The image analysis code is available at Ashrafi Lab’s GitHub repository. (https://github.com/ashrafilab/SynATP-Analysis)

## Notes

https://github.com/ashrafilab/SynATP-Analysis

### Competing Interest Statement

The authors have declared no competing interest.

### Summary of Updates

Updated graphs and detailed the analytic approach

